# Nano-particles from *Chaetomium* against Rice Blast

**DOI:** 10.1101/339283

**Authors:** JiaoJiao Song, Kasem Soytong

## Abstract

The effective isolate of *Chaetomium* sp actively against *Magnaporthe sp* causing rice blast were tested. Morphology and phylogenic identification was also confirmed *Chaetomium.sp. Magnporthe* sp proved to be virulent isolate to cause blast of rice. Bi-culture test proved that *Chaetomium sp* can be suppressed the growth of *Magnaporthe* sp. The fungal metabolites extracted from *Chaetomium* sp expressed antifungal *Magnaporthe* sp. Nano-particles derived from *Chaetomium* sp extracts exhibited significantly antifungal activity against *Magnaporthe* sp. Further investigation is to formulate the nanoparticles from active compounds of *Chaetomium* sp for plant immunity and to be applied in the rice fields.

**IMPORTANCE:** Rice blast is an economic important disease which caused by *Magnaporthe.* It causes yield loss and economic damage wherever rice is grown in Asia. It becomes the main disease due to wide distribution and most infection under favorable condition. The farmers have been applied several kinds of chemical fungicides that leading to the pathogen become resistant to those fungicides, and causes environmental pollution which the toxic chemicals residue in soil, water and surrounding environment. The significance of this research is to discover a new agricultural input as nanoparticles from *Chaetomium* to control the blast pathogen which will develop the nanoproduct to elicit immunity in rice, environmental protection and food safety.

## INTRODUCTION

Rice blast caused by *Magnaporthe oryzae* (Anamorph; *Pyricularia oryzae).* It is one of the first recorded as rice fever disease in China as early as 1637. Rice blast expanded through Asia, Latin America, and Africa and is now found in over 85 countries worldwide (1, 2). *M. oryzae* infects all parts of plants causing yield losses leading to shortage in many developing countries in recent years (1). Valent (3) stated that disease is epidemic in all continents wherever rice is grown and yield loss due to blast can be as high as 50% (4). The search for new effective disease control is needed to reduce or eliminate the chemical fungicides which causes the environment problem. Nanotechnology in agriculture becomes a new method to build up and restructure the materials at molecular level. Molecular nanotechnology involves constructing organic materials into defined structures, atom by atom or molecule by molecule (5). The application of nanotechnology in agriculture have increased attention in recent years (6). Researchers have actively investigated the synthesis of organic nanomaterials including different kinds of nanoparticle for biological properties (7, 8). The application of nanotechnology in agriculture are being explored (5) and could be helped the solution of overuse toxic chemicals faced by agriculture (9). Nanoparticles contain bioactive substances from natural products that can enable fast effective penetration through plant cuticles and tissues (10). It can provide efficient pest management in agriculture (11) and formulate to contain pesticides in colloidal suspension or powder to apply in agriculture (9), and increase the stability of active compounds (10). Application of natural products from *Chaetomium* spp. has been proven to be antifungal activity against several plant pathogens (12). Tann and Soytong (13) reported that nano particles constructed from *Chaetomium globosum* KMITL-N0805 expressed antifungal activity against *Curvularia lunata*, leaf spot of rice var. Sen Pidoa, and the tested nanoparticles can be controlled leaf spot of rice at 60 days. Furthermore, Tann and Soytong (14) reported nano-products derived from *Chaetomium cupreum* can be applied to reduce the incidence of rice leaf spot. This research was conducted to evaluate nano-particles constructed from *Chaetomium cochliodes* CTh05 for biological activity against *Magnaporthe oryzae* causing rice blast

## RESULTS

### Isolate of rice blast pathogen and *Chaetomium* sp

The symptoms of blast was isolated and morphological identified as *Magnaporthe.* Culture on PDA grown to full plate (9 cm) in10 days. Mycelia are septated and hyaline, producing conidiophores and 3 cells conidia. *Chaetomium.* from previous experiment (15), it was cultured on PDA for 3 weeks and shows olive-green to brown, producing perithecia and subglobose asci, one ascus has eight ascospores.

Molecular phylogenic identification was confirmed the species.

### Bi-culture Test

*Magnaporthe* sp. was inhibited by *Chaetomium* sp. The colony diameter of the antagonist averaged of 4.35 cm while the control was 9.00 cm, the colony growth inhibition was 51.66 *%* in 10 days. Thereafter incubation period was extended to 30 days and the colony inhibition averaged over 90 %.

### Bioactivity test of fungal metabolites from

*Chaetomium sp.* Result showed that CCoE gave significantly highest spore inhibition of 88 %, followed by CCoM and CCoH with spore inhibition of 80.74 and 67.57 %, respectively at 1,000 ppm in 12 days. The fungal metabolites CCoE, CCoM and CCoH showed antifungal activity against *Magnaporthe* sp. for spore inhibition (Table 1). Interestingly, at 15 ppm nanoCCoH, nanoCCoE and nanoCCoM gave significantly inhibited spore production. NanoCCoE gave significantly inhibited spore production of 68 % while nanoCCoM and nanoCCoH at 47 and 34 %, respectively in 12 days. The nanoparticles, nanoCCoE, nanoCCoM and nanoCCoH showed effective antifungal activity against *Magnaporthe* sp. It observed that the spores were abnormally shape and broken cells after treated with nanoparticles of *Chaetomium* sp and spores were normal shape in the non-treated control.

**Table 1.**
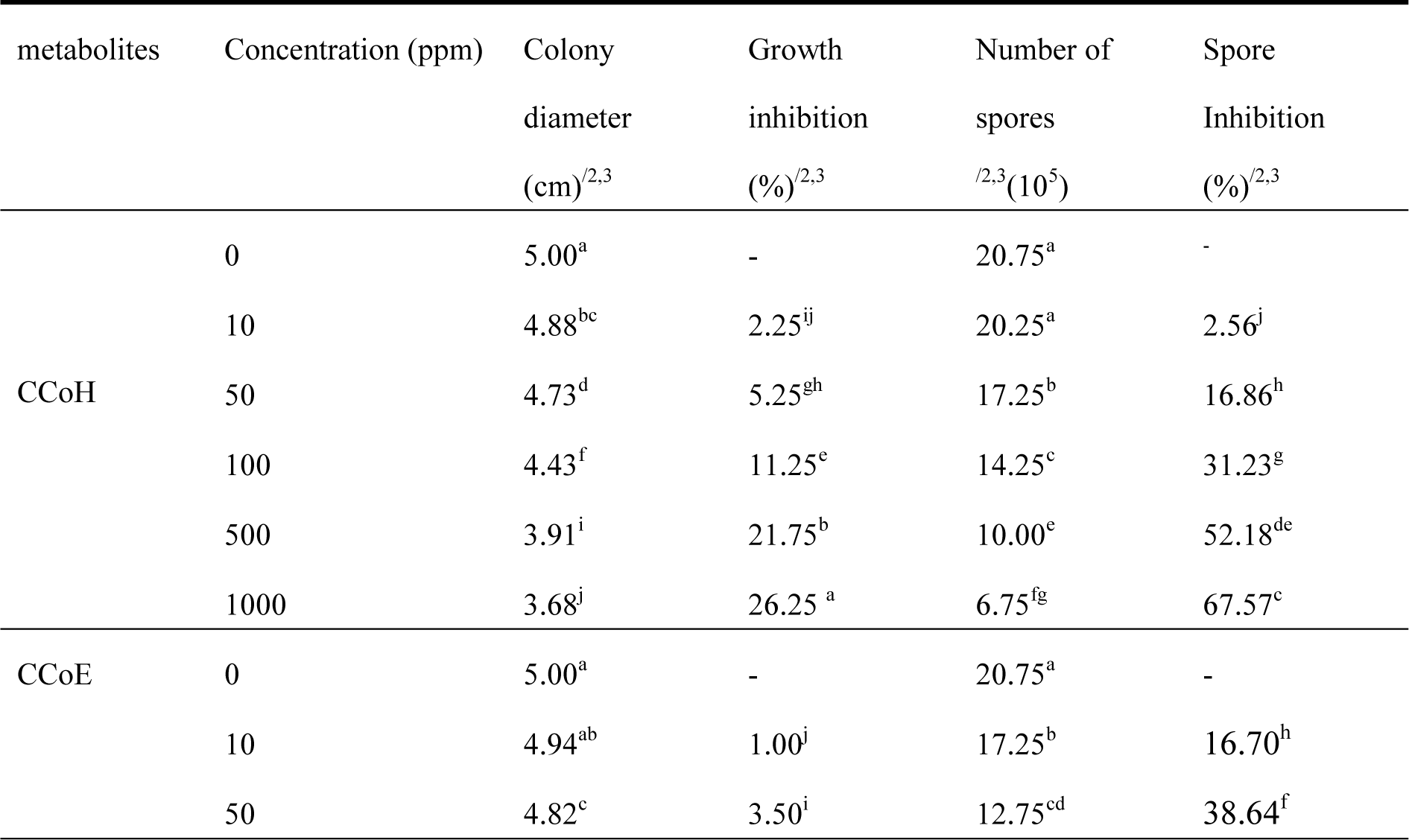
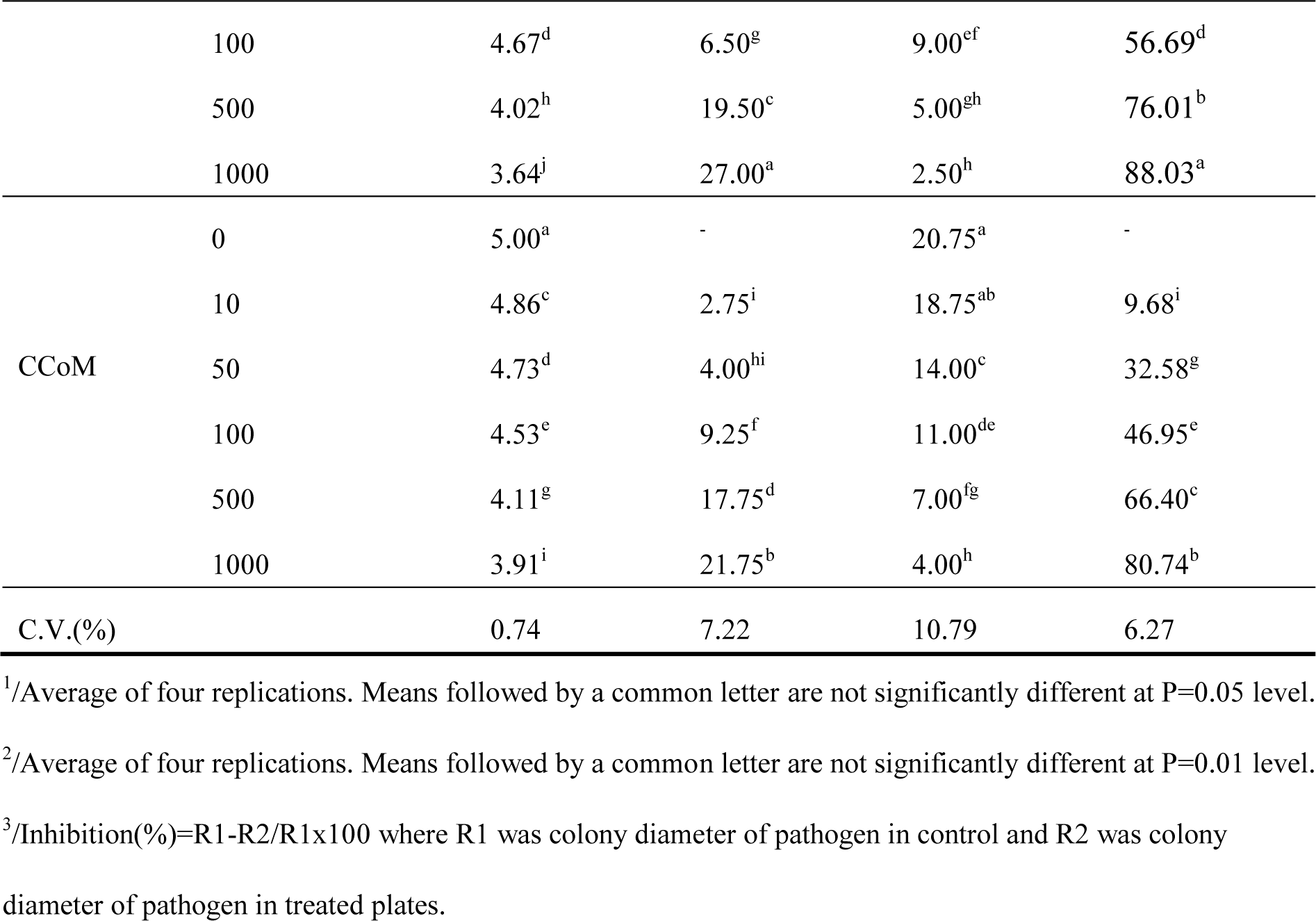
Fungal metabolites of *Chaetomium cochliodes* CTh05 against *Magnaporthe oryzae* PO1 at 12 days

## DISCUSSION

The symptoms of blast was isolated and confirmed morphological and phylogenic identified as *Magnaporthe oryzae. Magnaporthe* sp. proved to be aggressive isolate to cause blast of rice var. RD57. *Chaetomium* sp was confirmed by morphology and phylogenic identification and it was actively effective against *Magnaporthe* sp. causing rice blast. Bi-culture test proved that *Chaetomium sp* can suppress the growth of *Magnaporthe* sp. Previous report of Soytong (16) stated that *C. cochliodes* proved to be antagonized *Drechslera oryzae* (brown leaf spot of rice var Pittsanulok 2). It significantly inhibited colony growth and spores of tested pathogen in bi-culture test. Similar report stated that *Chaetomium cupreum* CC3003 inhibited *C. -lunata* (leaf spot of rice) in bi-culture test (14). The fungal metabolites of *Chaetomium* sp (CCoE, CCoM and CCoH) expressed antifungal activity against *Magnaporthe* sp. to inhibit spore production which the ED_50_ of 85, 144, 374 ppm. Soytong (16) reported that metabolites from *C. cochliodes* gave significant inhibition on the spore production of *D. oryzae* (brown leaf spot of rice) that the ED50 value was 66.45 ppm. Tann and Soytong (14) reported that crude hexane, EtOAc and MeOH extracts of *C. cupreum* C3003 inhibited spore production of *C. lunata* (leaf spot of rice) with the ED_50_ values of 6.41, 0.83 and 7.81 ppm, respectively.

The secondary metabolites from *Chaetomium* sp had been reported as the azaphilone types, cochliodones C and D, chaetoviridines E and F, *epi*-chaetoviridine A, and chaetoviridine A. The chaetoviridines E showed antimalarial activity against *Plasmodium falciparum.* Whereas, cochliodones C, chaetoviridines E and F expressed antimycobacterial activity against *Mycobacterium tuberculosis.* Moreover, chaetoviridines E and F exhibited cytotoxicity against the KB, BC1, and NCI-H187 cell lines (15). This research finding the fungal metabolites from *Chaetomium* sp actively against *M. oryzae* P01 causing rice blast pathogen.

Nano-particles derived from *Chaetomium* sp 5 (nanoCCoE, nanoCCoM and nanoCCoH) showed significant antifungal activity against *Magnaporthe* sp. with ED_50_ values of 9, 16 and 33 ppm, respectively. Similar report of Tann and Soytong (13) stated that nano-CGH, nano-CGE, and nano-CGM derived from *Chaetomium globosum* KMITL-N0805 showed antifungal activity against *Curvularia lunata* (leaf spot of rice) with ED_50_ values of 1.21, 1.19, and 1.93 ppm, respectively.

Dar and Soytong (17) stated that the fungal metabolites from *Chaetoimum globosum* and *Chaetoimum cupreum* which were reacted with polylactic acid and electropun at 25-30 kv, nanomaterial from *C. globosum* was 241 nanometers,and *C. cupreum* was 171 nanometers. Furthermore, the research finding showed abnormal spores of *Magnaporthe* sp.after treated with nanoparticles from *Chaetomium* sp. which was also reported by Tann and Soytong (13) that nanoparticles from *C. globosum* can be disrupted and distorted the pathogen cells, and loss of pathogenicity. With this, Singh *et al.* (18) stated that nanotechnology in agriculture is revolutionized by innovating new techniques to control diseases. In this research findings revealed that nano-particles constructed from fungal metabolites gave the ability for plant disease control. As said by Sharon et al. (19) that nanotechnology is helped in agriculture and reduced environmental pollution by applying nano particles with the ability to increase the efficiency of pesticide application at lower dose.

## MATERIALS AND METHODS

### Isolation of pathogen and pathogenicity test

The blast specimens were collected from blast symptom on leaves of rice var. RD 57. Isolation was done using tissue transplanting technique. Pure culture was maintained in rice flour agar (RFA) medium. Morphological identification was done under binocular compound microscope and followed the work of Ou (20). The pathogen is deposited at culture collection, Biocontrol Research Unit, Faculty of Agricultural Technology, KMITL, Bangkok, Thailand.

### Antagonistic fungus

*Chaetomium sp* used in this study which is deposited at culture collection, Biocontrol Research Unit, Faculty of Agricultural Technology, KMITL, Bangkok, Thailand. that reported to be produced antimicrobial activity against malaria (*Plasmopara falciparum*) and tuberculosis (*Mycobacterium tuberculosis)* as well as cancer cell lines (15). This isolate was cultured and incubated in potato dextrose agar (PDA) at room temperature (27-30 C), and morphological identified according to von Arx (21) and Soytong (22).

### Molecular identification

The genomic DNA were separately extracted from *Magnaporthe* and *Chaetomium*. Each fungus was cultured in potato dextrose broth (PDB) media for 3 days. The mycelia were freeze-dried, genomic DNA was extracted by modified CTAB (Cetyl trimethyl ammonium bromide) method, cleaned with 25mM EDTA by centrifugation. 100 mg of mycelia were crushed in liquid nitrogen. The fungal cells were lysed in CTAB buffer, β-mercaptoethanol, then incubated at 65°C for 1h in mixing tubes every 15 min. The lysate were extracted with an equal volume of chloroform/isoamyl alcohol (24:1), then centrifuged at 14,000 rpm for 5 min at 4°C. The aqueous phase was added 2μl Rnase (20μg/ml), and incubated 30 min at 37°C, then mixed with 50μl 10% CTAB. The genomic DNA was precipitated in isopropanol and centrifuged at 4°C for 20 min at 14,000 rpm. The resulting pellets were washed twice with 70% and 95% ethanol, air dried and dissolved in 100μl TE buffer at 37°C, overnight. The quality and quantity of extracted DNA were monitored by electrophoresis in a 1% agarose gel. Quantification was performed with comparison to the known dilution of lambda phage DNA. Polymerase chain reaction (PCR) was performed by ITS ribosomal DNA regions which amplified by PCR using the universal primers, ITS1 (5’-TCCGTAGGTGAACCTGCGG-3’) and ITS4 (5’-TCCTCCGCTTATTGATATGC-3’) (23). The 25μl reaction mixture contained 2.5μl 10 x PCR buffer, 0.625μl each dNTP (1.25mM), 0.5μl MgCl_2_, 1μl of each primer (20pmol/μl), 2 ng of DNA and 0.2μl of Taq DNA polymerase (1 U) were performed. PCR condition for the ITS regions were programed with initial denaturation at 95°C for 5 min, followed by 35 cycles of 95°C for 1 min., 56°C for 1 min., 72°C for 2 min., and a final extension at 72°C for 5 min. The amplified products (5μl) were visualized on 1% (w/v) agarose gel for the presence of a single amplified band. The amplified products were sequenced and aligned with comparison to the sequences in the GenBank by basic local alignment search tool (BLAST) analysis (24) at National Center for Biotechnology Information (NCBI) databases. The sequences of closely related organisms were downloaded to construct the phylogenetic trees, aligned through CLUSTALW using MEGA version 6.0 software (25) and a phylogenetic tree was designed by neighbor-joining method using the same software.

### Bi-culture test

The test was done by followed the method of Soytong and Quimio (26). *Chaetomium* and rice blast pathogen were transferred each 0.3 m culture agar plug to PDA at opposite sites. *Chaetomium* and rice blast pathogen were separately cultured where each isolate on PDA served as the controls. All plates were incubated at room temperature for 30 days. Data were collected as colony diameter (cm) and sporulation and subjected to analysis of variance and compared using Duncan Multiple’s Range Test (DMRT) at P=0.05 and 0.01.

### Fungal metabolites from *Chaetomium*

Biomass culture of *Chaetomium* was extracted with hexane, ethyl acetate and methanol using method described Phonkerd *et al.* (15) to give crude hexane (CCoH), ethyl acetate (CCoE) and methanol (CCoM) extracts. The experiment was set up as two factor factorial experiment in CRD with four replications. Factor A represented crude extracts and factor B represented various concentrations of 0, 10, 50, 100, 500 and 1,000 ppm. The culture agar plug of 3 mm was transferred to the middle PDA plate in each treatment, then incubated at room temperature (27-30 C) for 15 days. Data were collected as colony diameter (cm) and counted number of spores using haemacytometer. Data were statistically computed analysis of variance (ANOVA). Treatment means were compared using DMRT at P=0.05 and 0.01. The effective dose (ED50) was computed using probit analysis program.

### Nano-particles

The crude extracts from *Chaetomium* were separately incorporated into polylactic acid (PLA)-based nanoparticles through electrospinning by followed the method of Dar and Soytong (17) to yield nanoCCoH, nanoCCoM and nanoCCoE, respectively. Nano CCoH, CCoM and CCoE were tested to inhibit the rice blast pathogen using two factorial in CRD with four replications. Factor A was nanoparticles, and factor B was 0, 3, 5, 7, 10 and 15 ppm. CRD was designed for the experiment which repeated four times. Data were collected as colony diameter (cm) and counted number of spores using haemacytometer, and statistically calculated analysis of variance (ANOVA). Means were compared using DMRT at P=0.05 and 0.01. The effective dose (ED50) was calculate using probit analysis program.

## ACKNOWLEDGMENT

I would like to acknowledge the King Mongkut’s Institute of Technology Ladkrabang (KMITL) to offer a PhD Scholarship and research fund is supported by Faculty of Agricultural Technology, KMITL and further research fund [KREF126004] supported by KMITL, Bangkok, Thailand. The financial support from Thailand Research Fund (Grant No RTA5980002) is also gratefully acknowledged.

